# Alzheimer’s-like remodeling of neuronal ryanodine receptor in COVID-19

**DOI:** 10.1101/2021.02.18.431811

**Authors:** Steve Reiken, Haikel Dridi, Leah Sittenfeld, Xiaoping Liu, Andrew R Marks

## Abstract

COVID-19, caused by SARS-CoV-2 involves multiple organs including cardiovascular, pulmonary and central nervous system. Understanding how SARS-CoV-2 infection afflicts diverse organ systems remains challenging^1,2^. Particularly vexing has been the problem posed by persistent organ dysfunction known as “long COVID,” which includes cognitive impairment^3^. Here we provide evidence linking SARS-CoV-2 infection to activation of TGF-ß signaling and oxidative overload. One consequence is oxidation of the ryanodine receptor/calcium (Ca^2+^) release channels (RyR) on the endo/sarcoplasmic (ER/SR) reticuli in heart, lung and brains of patients who succumbed to COVID-19. This depletes the channels of the stabilizing subunit calstabin2 causing them to leak Ca^2+^ which can promote heart failure^4,5^, pulmonary insufficiency ^6^ and cognitive and behavioral defects^7–9^. *Ex-vivo* treatment of heart, lung, and brain tissues from COVID-19 patients using a Rycal drug (ARM210)^10^ prevented calstabin2 loss and fixed the channel leak. Of particular interest is that neuropathological pathways activated downstream of leaky RyR2 channels in Alzheimer’s Disease (AD) patients were activated in COVID-19 patients. Thus, leaky RyR2 Ca^2+^ channels may play a role in COVID-19 pathophysiology and could be a therapeutic target for amelioration of some comorbidities associated with SARS-CoV-2 infection.

## Main

Patients suffering from COVID-19 exhibit multi-organ system failure involving not only pulmonary ^1^ but also cardiovascular ^2^, neural ^11^ and other systems. The pleiotropy and complexity of the organ system failures both complicate the care of COVID-19 patients, and contribute to a great extent to the morbidity and mortality of the pandemic^12^. Clinical data^13^ indicate that severe COVID-19 most commonly manifests as viral pneumonia-induced acute respiratory distress syndrome (ARDS). Respiratory failure results from severe inflammation in the lungs, which arises when COVID-19 infects lung cells and damages them. Cardiac manifestations are multifactorial, and include hypoxia, hypotension, enhanced inflammatory status, angiotensin-converting enzyme 2 (ACE2) receptor downregulation, endogenous catecholamine adrenergic status, and direct viral myocardial damage^14,15^. Moreover, patients with underlying cardiovascular disease or comorbidities, including congestive heart failure, hypertension, diabetes, and pulmonary diseases, are more susceptible to infection by SARS-CoV-2, with higher mortality^14,15^. In addition to respiratory and cardiac manifestation, it has been reported that 36.4% of patients with COVID-19 develop neurological symptoms, including headache, disturbed consciousness, and paresthesia ^16^. Brain tissue edema, stroke, partial neuronal degeneration in deceased patients, and neuronal encephalitis have also been reported^2,16–18^. Furthermore, another pair of frequent symptoms of infection by SARS-CoV-2 are hyposmia and hypogeusia, the loss of the ability to smell and taste, respectively^11,19^. Interestingly, hyposmia has been reported in early stage AD^11^ and Alzheimer type II astrocytosis has been observed in neuropathology studies of COVID-19 patients^18^. Despite the myriad of neurological symptoms, manifestations, and complications reported with SARS-CoV-2 infection^20,21^, a systematic study of COVID-19 brain neuropathology is still needed^22^.

Systemic failure in COVID-19 patients is likely due to SARS-CoV-2 invasion via the ACE2 receptor^17^, which is highly expressed in pericytes of human heart^23^, epithelial cells of the respiratory tract^24^, kidney, intestine, and blood vessels. ACE2 is also expressed in the brain, especially in the brainstem, specifically the respiratory center and hypothalamus, the thermal center, and cortex^25^, which renders these tissues more vulnerable to viral invasion. The primary consequences of SARS-CoV-2 infection are inflammatory responses and oxidative stress in multiple organs and tissues^26^–^28^. Recently it has been shown that the high neutrophil to lymphocyte ratio observed in critically ill patients with COVID-19 is associated with excessive levels of reactive oxygen species (ROS) and that ROS induced tissue damage, contributing to COVID-19 disease severity^26^. The impact of oxidative stress on calcium (Ca^2+^) homeostasis, a key determinant of cardiac function and rhythm^29^, as well as neuronal activity, remain to be elucidated.

Recent studies have explored the relationship between ACE2 and TGF-β, demonstrating the inverse relationship between the two. In cancer models, decreased levels of ACE2 correlated with increased levels of TGF-β^30^. In the context of SARS-CoV-2 infection, downregulation of ACE2 has been observed, leading to increased fibrosis formation, as well as upregulation of TGF-β and other inflammatory pathways^31^. Finally, patients with severe COVID-19 symptoms had higher blood serum TGF-β concentrations than those with mild symptoms^32^, thus further implicating the role of TGF-β in this disease and warranting further investigation on the topic.

Downregulation of ACE2 has been observed in SARS-CoV-2 infection, leading to increased fibrosis formation, as well as upregulation of TGF-β and other inflammatory pathways^31^. Interestingly, reduced angiotensin/ACE-2 activity has been associated with Tau hyper-phosphorylation and increased amyloid-β pathology in animal models of Alzheimer disease^33,34^. The link between reduced ACE2 activity and increased TGF-β and Tau signaling in the context of SARS-CoV-2 infection needs further exploration.

Our laboratory has shown that stress-induced ryanodine receptor (RyR)/intracellular calcium release channel post-translational modifications, including oxidation and protein kinase A (PKA) hyper-phosphorylation related to activation of the sympathetic nervous system and the resulting hyper-adrenergic state, deplete the channel stabilizing protein (calstabin) from the channel complex, destabilizing the closed state of the channel causing it to leak in multiple diseases^6–9,29,35,36^. Increased transforming growth factor-β (TGF-β) activity can lead to RyR modification and leaky channels^37^ and SR Ca^2+^ leak can cause mitochondrial Ca^2+^ overload and dysfunction^36^. Increased TGF-β activity and^38^ mitochondrial dysfunction^39^ are also associated with SARS-CoV-2 infection.

Here we show that SARS-CoV-2 infection is associated with oxidative stress and activation of the TGF-β signaling pathway in heart, lung and brain of patients who have succumbed to COVID-19. One consequence is RyR oxidation, particularly through NADPH oxidase 2 (NOX2), rendering the channels leaky to Ca^2+^ which may play a role in the pathological manifestations of SARS-CoV-2 infection in heart, lung and brain. Moreover, the hyper-adrenergic state observed in COVID-19 patients causes PKA hyper-phosphorylation of RyR2 on Serine 2808, loss of the stabilizing subunit calstabin2 from the channel complex and leaky RyR2 channels in heart, lung, and brain. We also demonstrate that SARS-CoV-2 infection activates biochemical pathways linked to the Tau pathology associated with AD and that leaky calcium channels may be a potential therapeutic target for the respiratory, cardiac and neuronal complications associated with COVID-19.

## Results

### Oxidative stress and TGF-β activation

Oxidative stress was determined in heart, lung, and brain tissues from COVID-19 patient autopsy tissues and controls by measuring the ratio of glutathione disulfide (GSSG) to glutathione (GSH). COVID-19 patients exhibited significant oxidative stress with a 7.7, 13.6, and 3.2-fold increase in GSSG/GSH ratios in heart, lung, and brain compared to controls respectively **(Figure 1A)**. In order to determine if SARS-CoV-2 infection also increases tissue TGF-β activity, we measured SMAD3 phosphorylation, a downstream signal of TGFβ, in control and COVID-19 tissue lysates **(Figure 1B)**. Phosphorylated SMAD3 (pSMAD3) levels were increased in COVID-19 heart, lung, and brain lysates compared to controls, indicating that SARS-CoV-2 infection increased TGF-β signaling in these tissues. Interestingly, brain tissues from COVID-19 patients exhibited activation of the TGF-β pathway, despite the absence of the detectable (by immunohistochemistry and PCR, data not shown) virus in these tissues. These results suggest that the TGF-β pathway is activated systemically by SARS-CoV-2 resulting in its upregulation in brain as well as other organs.

**Figure 1.**
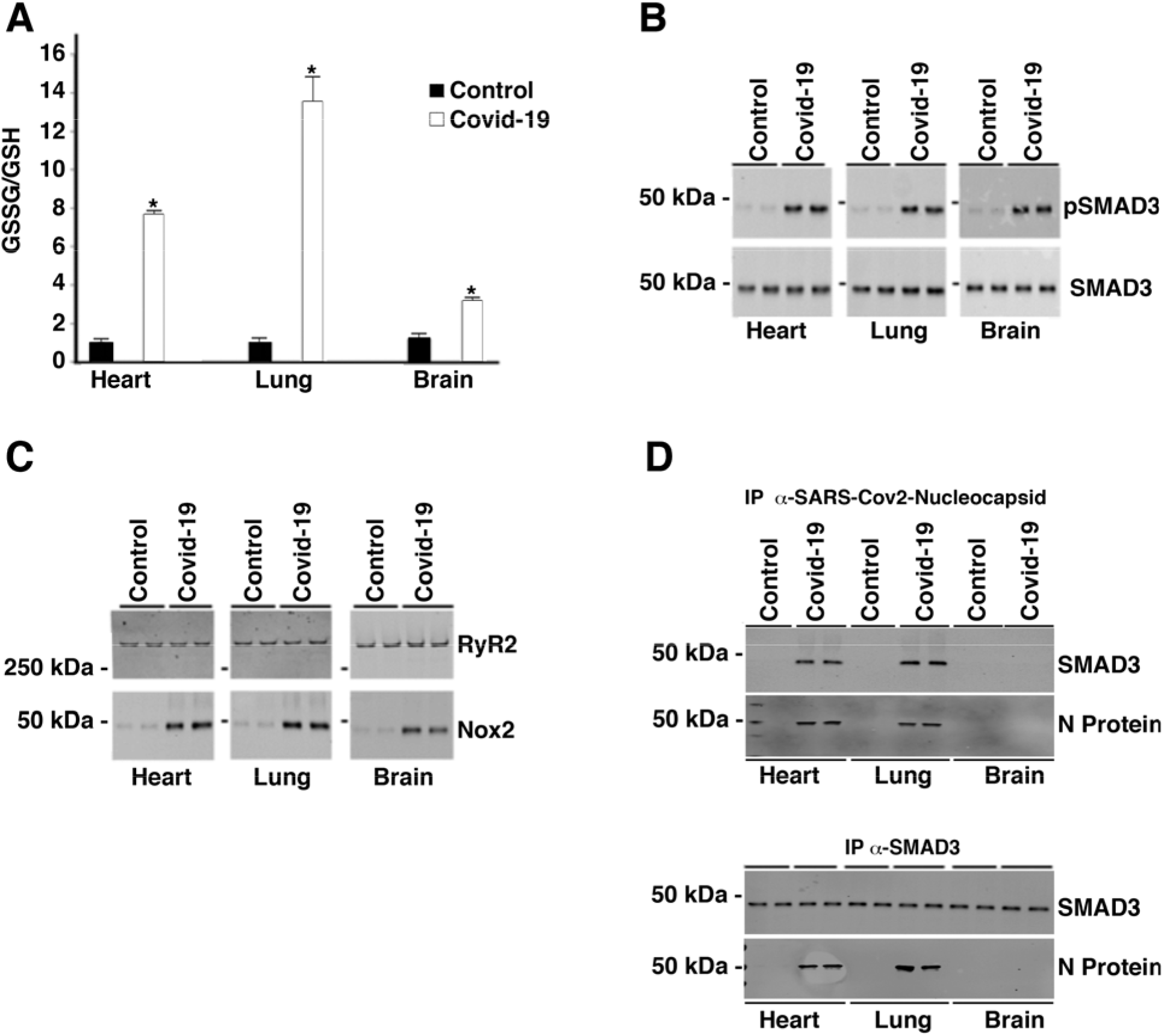
Increased oxidative stress in heart, lung, and brain of COVID-19 patients. **(A)** Bar graph depicting the GSSG/GSH ratio from control (n=2) and COVID-19 (n=2) tissue lysates. *p<0.05 control vs COVID-19. **(B)** Western blots showing Phospho-SMAD3 and total SMAD3 from control (n=2) and COVID-19 (n=2) tissue lysates. **(C).** Western blots showing the co-immunoprecipitation of NOX2 with RyR2 from control (n=2) and COVID-19 (n=2) tissue lysates. **(D).** Western blots showing the co-immunoprecipitation of SARS-CoV-2-nucleocapsid (N protein) with SMAD3. Upper panel are westerns of immunoblots blotted for both N protein and SMAD3 from lysates that were immunoprecipitated with anti-SARS-CoV-2-nucleocapsid antibody. Bottom panel are westerns of immunoblots blotted for both N protein and SMAD3 from lysates that were immunoprecipitated with anti-SMAD3 antibody.

### RyR2 channel oxidation and leak

RyR channels may be oxidized as a consequence of activation of the TGF-β signaling pathway^37^. NOX2 binding to RyR2 causes oxidation of the channel that activates the channel manifested as an increased open probability. When the oxidization of the channel is at pathologic levels there is destabilization of the closed state of the channel resulting in spontaneous Ca^2+^ release or leak^35,40^. To determine the effect of the increased TGF-β signaling associated with SARS-Cov-2 infection on NOX2/RyR2 interaction, RyR2 and NOX2 were coimmunoprecipitated from heart, lung, and brain lysates of COVID-19 patients and controls. NOX2 associated with RyR2 in heart, lung, and brain tissues from SARS-CoV-2 infected individuals was increased compared to controls **(Figure 1C)**.

The SMAD3 proteins involved in TGF-β signaling were shown to interact with the nucleocapsid (N) protein in SARS-CoV1^38^, interfering with healthy apoptotic pathways. As a result, apoptosis of SARS-CoV-2 infected host cells is blocked and formation of tissue fibrosis is promoted, especially in lung, thus contributing to the respiratory distress and subsequent pulmonary failure associated with the disease. SARS-CoV-2 infection may also activate TGF-β signaling through a direct interaction of the viral nucleocapsid protein (N protein) with SMAD3. This direct interaction is shown in **Figure 1D,** which demonstrates that SMAD3 and SARS-CoV-2 N protein co-immunoprecipitated from COVID-19 heart and lung lysates using antibodies specific for either protein. The brain lysates used in this study did not contain detectable (by immunoprecipitation/immunoblotting) SARS-CoV-2 N protein, indicating that indirect effects of the inflammatory/oxidative response may be responsible for the alterations in brain biochemistry observed in **Figure 1.**

Given the increased oxidative stress and increased NOX2 binding to RyR2 seen in COVID-19 tissues, RyR2 post-translational modifications in these tissues were investigated. Immunoprecipitated RyR2 from heart, lung, and brain lysates demonstrated increased oxidation, PKA phosphorylation on Serine 2808, and depletion of the stabilizing protein subunit calstabin2 in SARS-CoV-2 infected tissues compared to controls **(Figure 2A-C)**.

**Figure 2.**
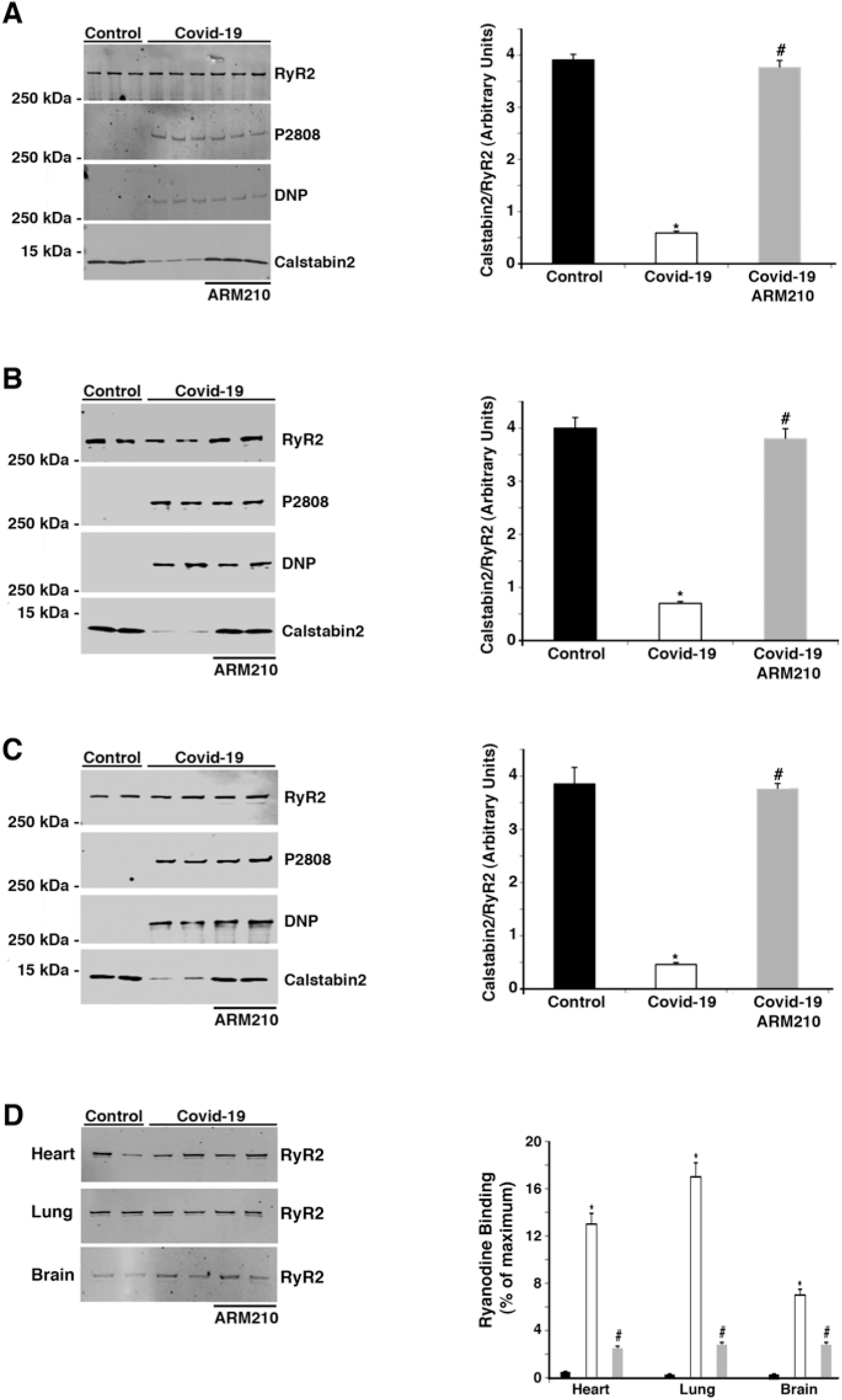
RyR2 post-translational modifications and calstabin2 binding in COVID-19 tissues. **(A)** Western blots (left, panel) and quantification of calstabin2/RyR2 association from heart lysates (right panel). **(B)** Western blots (left panel) and quantification of calstabin2/RyR2 association from lung lysates (right panel). **(C)** Western blots (left panel) and quantification of calstabin2/RyR2 association from brain lysates (right panel). **(D)** ^3^[H]ryanodine binding from immunoprecipited RyR2. Bar graphs show ryanodine binding at 150 nM Ca^2+^ as a percent of maximum binding (Ca2+ = 20 μM). Data are mean ± SD. t-test revealed *p<0.05 control vs COVID-19; # p<0.05 COVID-19 vs COVID-19+ARM210.

RyR channel activity was determined by binding of ^3^[H]ryanodine, which binds only to the open state of the channel. RyR2 was immunoprecipitated from tissue lysates and ryanodine binding to the immunoprecipitated was measured at both 150 nM and 20 μM free Ca^2+^. The total amount of RyR immunoprecipitated was the same for control and COVID-19 samples (immunoblots in Figure 2D). However, RyR2 channels from SARS-CoV-2 infected heart, lung, and brain tissue demonstrate abnormally high activity compared to channels from control tissues at physiologically resting conditions (150 nM free Ca^2+^), when channels should be closed (Figure 2D). This biochemical/functional remodeling of the channel results in what is known as “the biochemical signature” of leaky RyR2^4,41^ that destabilizes the closed state of the channel. This leads to SR/ER chronic Ca^2+^ leak which contributes to the pathophysiology of various diseases ^7,9,37,42^. Rebinding of calstabin2 to RyR2, using a Rycal, has been shown to reduce SR/ER Ca^2+^ leak, despite the persistence of the channel remodeling. Indeed, calstabin2 binding to RyR2 was increased when COVID-19 patient heart, lung and brain tissue lysates were treated *ex-vivo* with the Rycal drug ARM210 **(Figure 2A-C).** RyR2 activity at resting Ca^2+^ concentration was also decreased by Rycal treatment **(Figure 2D).**

### Activation of Alzheimer’s Disease linked signaling

Leaky ryanodine receptors have recently been implicated in the neurodegenerative processes that contribute to the etiology of Alzheimer’s and Huntington’s disease^7,8^. Abnormal Ca^2+^ handling can contribute to mitochondrial Ca^2+^ overload, dysregulation of Ca^2+^-dependent enzymes such as AMP-activated protein kinase (AMPK), cyclin-dependent kinase 5 (CDK5), and enhanced calpain activity^7^. Activation of these enzymes in response to elevated cytosolic Ca^2+^ levels is upstream of both Tau and amyloid deposits in Alzheimer’s disease brains, and could play an important role in the “brain fog” associated with COVID-19. Brain lysates from COVID-19 patients’ autopsies demonstrated increased AMPK and GSK3β phosphorylation compared to controls (**Figure 3A-B**). Activation of these kinases in SARS-CoV-2 infected brain leads to a hyper-phosphorylation of Tau similar to that observed in Alzheimer Tau pathology. COVID-19 brain lysates showed increased Tau phosphorylation at S199 and S202/T205 **(Figure 3A-B)**. We also observed an increased p25 expression, the neurotoxic activator of CDK5 **(Figure 3C-D)**. CDK5 plays an important role in amyloid precursor protein (APP) processing in AD. However, the increased p25/CDK5 observed in COVID-19 brain lysates did not activate the amyloid beta pathway as no differences observed in BACE1 or BCTF levels between control and COVID-19 brain lysates **(Figure 3C-D)**.

**Figure 3.**
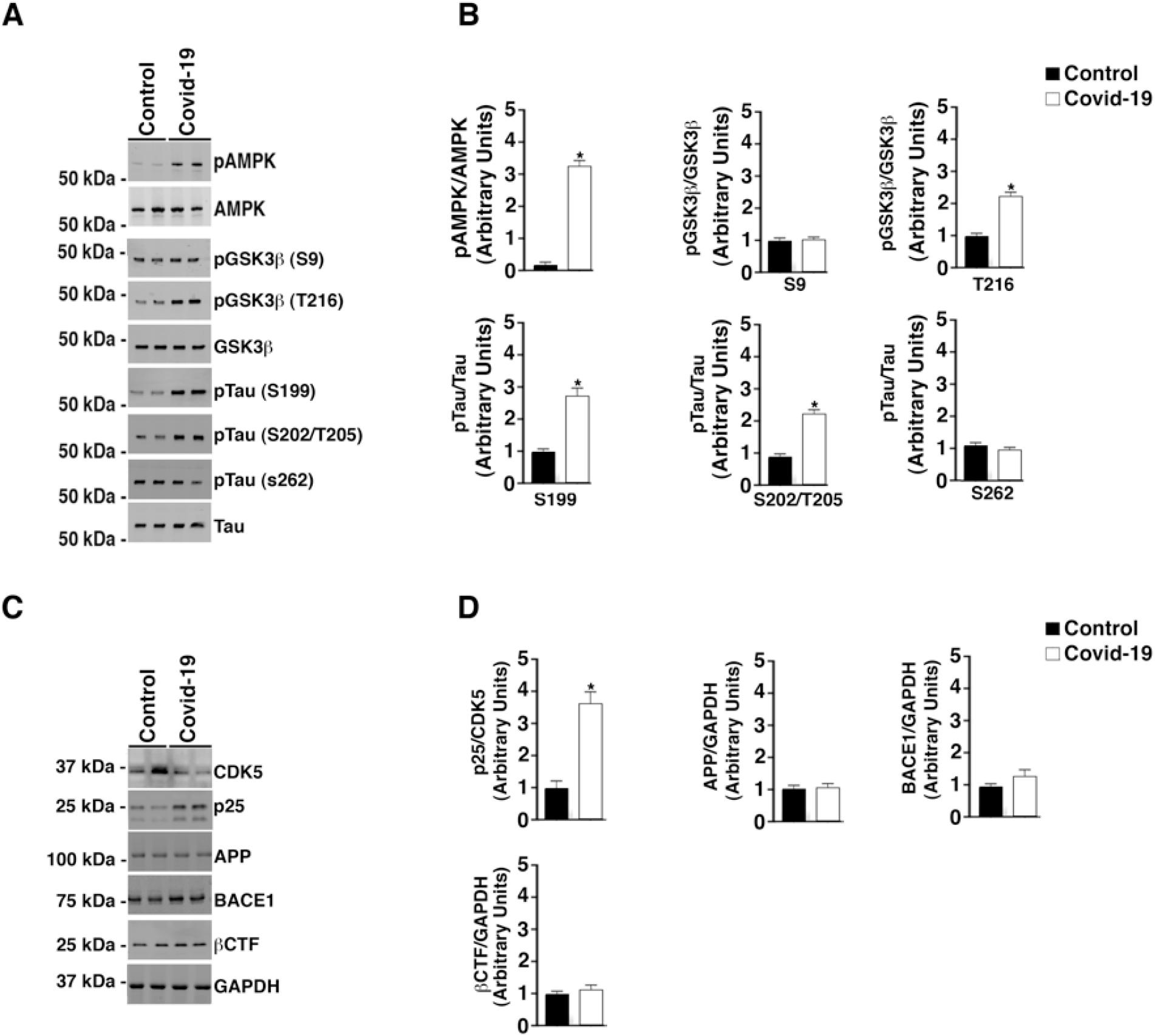
Hyperphosphorylation of Tau but normal APP processing in COVID-19 brain lysates. **(A)** Brain lysates were separated by 4-20% PAGE. Immunoblots were developed for pAMPK, AMPK, GSK3β, pGSK3β (S9, T216), Tau, pTau (S199, S202/T205, S262). **(B)** Bar graphs showing quantification of phosphorylated AMPK, pGSK3β and pTau from westerns. (**C**). immunoblots developed for CDK5, CDK5 truncated activator p25, APP, BACE1, and β CTF. **(D)** Bar graphs showing quantification p25/CDK5 and expression of APP, BACE1, and β CTF compared to loading control (GAPDH). Data are mean ± SD. t-test revealed *p<0.05 control vs COVID-19.

## Discussion

The molecular basis of how SARS-CoV-2 infection affects various tissues is not well understood, and questions regarding the role of defective Ca^2+^ signaling in heart, lung, and brain remain unanswered. In this study, we propose a potential mechanism that may contribute to systemic organ failure caused by SARS-CoV-2: defective Ca^2+^ regulation and its downstream signaling.

TGF-β belongs to a family of cytokines involved in the formation of cellular fibrosis by promoting epithelial-to-mesenchymal transition, fibroblast proliferation, and differentiation^43^. TGF-β activation has been shown to induce fibrosis in lung and other organs by activation of the SMAD-dependent pathway. We have previously reported that TGF-β/SMAD3 activation leads to NOX2/4 translocation to the cytosol and association with RyR channels promoting oxidization of the channels and depletion of the stabilizing subunit calstabin in skeletal muscle and in heart ^35,37^ Alteration of Ca^2+^ signaling may be particularly crucial in COVID-19 infected patients with cardiovascular/neurological diseases due, in part, to the multifactorial RyR2 remodeling following the cytokine storm, increased TGF-β activation, and increased oxidative stress. Moreover, SARS-CoV-2 infected patients exhibited increased endogenous adrenergic state, which leads to hyper-phosphorylation of RyR2 channels, as observed in this study, promoting pathological remodeling of the channel and exacerbating defective Ca^2+^ regulation that likely result in cardiomyopathies and arrhythmias. In line with our observation, a recent study has reported evidence of takotsubo cardiomyopathy caused by a stroke and an acute COVID-19 induced sympathetic stimulation and catecholamines surge which has led to death^44^.

## Conclusion

Our data indicate a role for leaky RyR2 in the pathophysiology of SARS-CoV-2 infection (**Figure 4**). We show increased systemic oxidative stress and activation of the TGF-β signaling pathway in lung, heart and brain of COVID-19 patients that correlates with oxidation-driven biochemical remodeling of the ryanodine receptor calcium release channel (RyR2). This RyR2 remodeling results in intracellular calcium leak which can promote heart failure, pulmonary insufficiency and cognitive dysfunction. Of particular interest is that leaky RyR2 channels in the brain were associated with activation of neuropathological pathways that are also found in the brains of Alzheimer’s Disease patients. *Ex-vivo* treatment of COVID-19 lung, heart and brain using a Rycal drug (ARM210) that targets RyR2 channels prevented intracellular Ca^2+^ leak in patient samples.

**Figure 4.**
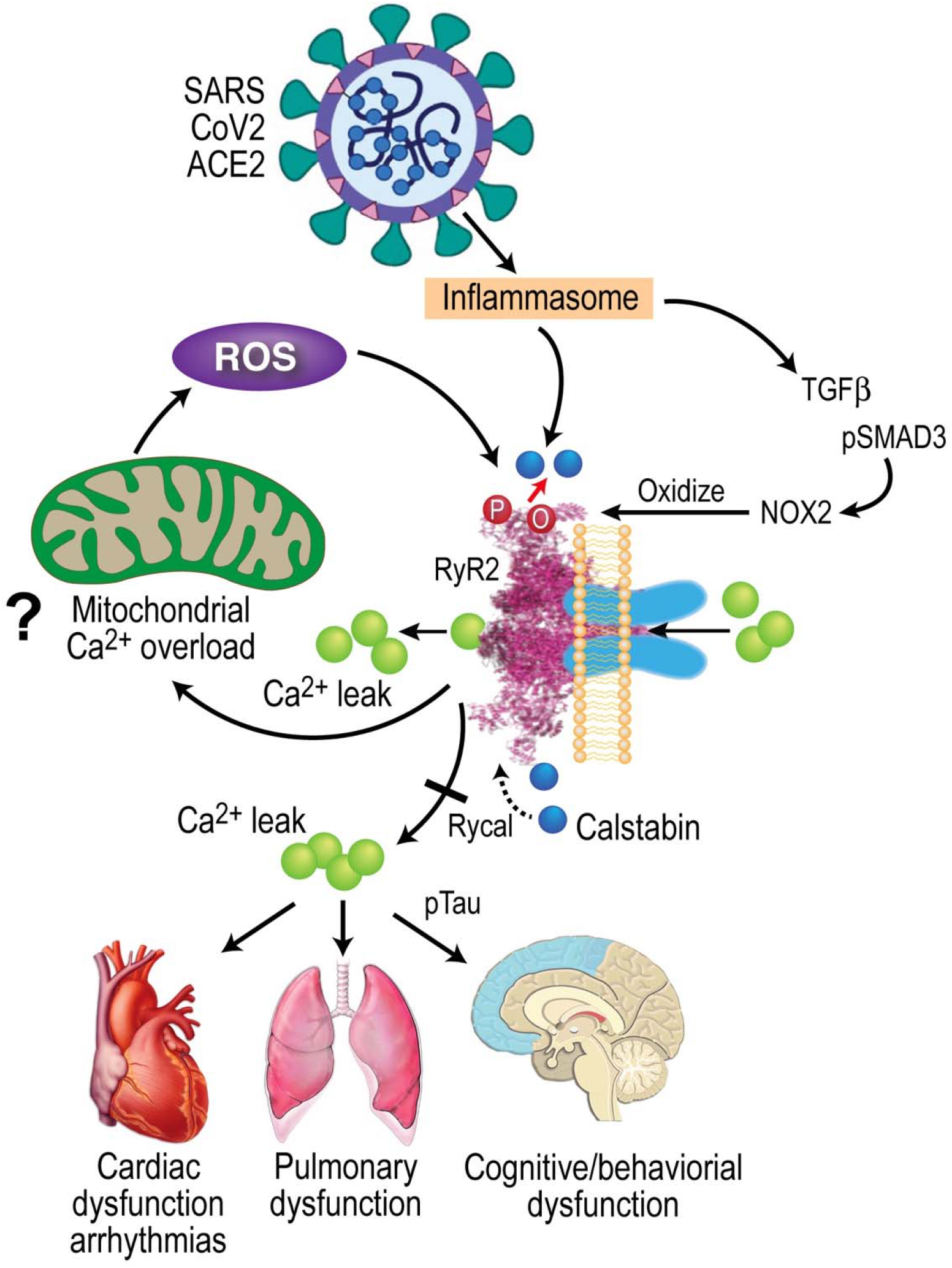
SARS-CoV-2 infection results in leaky RyR2 that may contribute to cardiac, pulmonary, and cognitive dysfunction. SARS-CoV-2 infection targets cells via the ACE2 receptor inducing inflammasome stress response/activation of stress signaling pathways. This results in increased TGF-β signaling, which activates SMAD3 (phosphoSMAD3, pSMAD), this increases NOX2 expression and the amount of NOX2 associated with RyR2. Increased NOX2 activity at RyR2 oxidizes the channel causing calstabin depletion from the channel macromolecular complex, destabilization of the closed state, and ER/SR calcium leak that is known to contributes to cardiac dysfunction^4,5^, arrhythmias^28^, pulmonary insufficiency^6,8^, and cognitive and behavioral abnormalities associated with neurodegenreation^7,8^. Rycal drugs fix the RyR channel leak by restoring calstabin binding and stabilizing the channel closed state. Fixing leaky RyR may improve cardiac, pulmonary and neural function in COVID-19.

## Acknowledgments

This work was supported by grants from the NIH (R01HL145473, R01DK118240, R01HL142903, R01HL140934, R01AR070194, T32HL120826)

## Author contributions

HD, LS, SR, XL and ARM designed experiments, analyzed data and edited/wrote the paper.

## Data availability

The data supporting the findings of this are documented within the paper and are available from the corresponding author upon request.

## Methods

### Human samples

De-identified human heart, lung, and brain tissue from the Covid BioBank at Columbia University. The Columbia University biobank functions under standard operating procedures, quality assurance, and quality control for sample collection and maintenance. Age and gender-matched controls exhibited absence of neurological disorders and cardiovascular or pulmonary diseases.

### Lysate preparation and western blots

Tissue (50 mg) were isotonically lysed using a dounce homogenizer in 0.25 ml of 10 mM Tris Maleate (pH 7.0) buffer plus protease inhibitors (Complete inhibitors from Roche). Samples were centrifuged at 8,000 x g and the protein concentration of the supernatants were determined by Bradford assay. To determine protein levels in tissue lysates, tissue proteins (20 μg) were separated by 4-20% SDS-PAGE and immunoblots were developed using antibodies against pSMAD3 (Abcam, 1:1000), SMAD3 (Abcam, 1:1000), AMPK (Abcam, 1:1000), pAMPK (Abcam, 1:1000), CDK5 (Thermofisher, 1: 1,000), and p25 (Thermofisher, 1:1000). Tau (Thermofisher, 1:1000), P-Tau (S199, Thermofisher, 1:1000), P-Tau (S202/T205, Abcam, P-Tau (S262, Abcam, 1:1000). 1:1000), GSK3β (Abcam, 1: 2,000)), p-GSK3β (S9, Abcam, 1: 2,000), p-GSK3β (T216, Abcam, 1: 2,000), APP (Abcam, 1: 2,000), BACE1 (Abcam, 1: 2,000), GAPDH (Santa Cruz Bioteck, 1:1000), βCTF (Abcam, 1: 2,000), SARS Cov-2 nucleocapsid (Thermofisher, 1: 1,000).

### Analysis of ryanodine receptor complex

Tissue lysates (0.1 mg) were treated with buffer or 10 μM Rycal (ARM210) at 4°C. RyR2 were immunoprecipitated from 0.1 mg lung, heart and brain using an anti-RyR2 specific antibody (2 μg) in 0.5 ml of a modified radioimmune precipitation assay buffer (50 mm Tris-HCl, pH 7.2, 0.9% NaCl, 5.0 mm NaF, 1.0 mm Na3VO4, 1% Triton X-100, and protease inhibitors) overnight at 4 °C. RyR2 specific antibody was an affinity-purified polyclonal rabbit antibody using the peptide CKPEFNNHKDYAQEK corresponding to amino acids 1367-1380 of mouse RyR2 with a cysteine residue added to the amino terminus. The immune complexes were incubated with protein A-Sepharose beads (Sigma) at 4 °C for 1 h, and the beads were washed three times with radioimmune precipitation assay buffer. The immunoprecipitates were size-fractionated on SDS-PAGE gels (4-20 % for RyR2, calstabin and NOX2) and transferred onto nitrocellulose membranes for 2 h at 200 mA. Immunoblots were developed using the following primary antibodies: anti-RyR2 (Affinity Bioreagents, 1:2000), anti-phospho-RyR-Ser(P)-2808 (Affinity Bioreagents 1:5000), anti-calstabin (FKBP12 C-19, 1:1000, Santa Cruz Biotechnology, Inc., Santa Cruz, CA), anti NOX2 (Abcam, 1:1000). To determine channel oxidation, the carbonyl groups in the protein side chains were derivatized to DNP by reaction with 2,4-dinitrophenylhydrazine. The DNP signal associated with RyR was determined using a specific anti-DNP antibody according to the manufacturer’s instructions (Millipore, Billerica, MA). All immunoblots were developed and quantified using an Odyssey system (LI-COR Biosciences, Lincoln, NE) with IR-labeled anti-mouse and anti-rabbit IgG (1: 10,000 dilution) secondary antibodies.

### Ryanodine Binding

RyR2 were immunoprecipitated from 1.5 mg of tissue lysate using an anti-RyR2 specific antibody (25 μg) in 1.0 ml of a modified RIPA buffer overnight at 4 °C. The immune complexes were incubated with protein A-Sepharose beads (Sigma) at 4 °C for 1 h, and the beads were washed three times with RIPA buffer, followed by 2 washes with ryanodine binding buffer (10 mM Tris-HCl, pH 6.8, 1 M NaCl, 1 % CHAPS, 5 mg/ml phosphatidylcholine, and protease inhibitors). Immunoprecipitates were incubated in 0.2 ml of binding buffer containing 20 nM [^3^H] ryanodine and either of 150 nM and 20 μm free Ca^2+^ for 1 h at 37 °C. Samples were diluted with 1 ml of ice-cold washing buffer (25 mm Hepes, pH 7.1, 0.25 m KCl) and filtered through Whatman GF/B membrane filters pre-soaked with 1% polyethyleneimine in washing buffer. Filters were washed three times with 5ml of washing buffer. The radioactivity remaining on the filters is determined by liquid scintillation counting to obtain bound [^3^H] ryanodine. Nonspecific binding was determined in the presence of 1000-fold excess of non-labeled ryanodine.

### GSSH/GSH ratio measurement and SMAD3 phosphorylation

Approximately 20 mg of suspended in 200 μL of ice-cold PBS/0.5% NP-40, pH6.0 for lysis. Tissue is homogenized with a Dounce homogenizer with 10 – 15 passes. Samples are centrifuged at 8,000 x g for 15 minutes at 4°C to remove any insoluble material. Supernatant is transferred to a clean tube. Deproteinizing of the samples is accomplished by adding 1 volume ice cold 100% (w/v) TCA into 5 volumes of sample and vortexing briefly to mix well. After incubating for 5 min on ice, samples are centrifuged at 12,000 x g for 5 minutes at 4°C and the supernatant is transferred to a fresh tube. The samples are neutralized by adding NaHCO3 to supernatant and vortexing briefly. Samples are centrifuged at 13,000 x g for 15 minutes at 4°C and supernatant is collected. Samples are now deproteinized, neutralized, and TCA has been removed. The samples are ready to use in the assay. The GSSG/GSH is determined using the ratio detection assay kit (Abcam, ab 138881). Briefly, in two separate assay reactions, GSH (reduced) is measured directly with a GSH standard and Total GSH (GSH + GSSG) is measured by using a GSSG standard. A 96 well plate is set up with 50 μL duplicate samples and standards with known concentrations of GSH and GSSG. A Thiol green indicator is added and the plate is incubated for 60 min at room temperature. Fluorescence at Ex/Em = 490/520 nm is measured with a fluorescence microplate reader, and the GSSG/GSH for samples are determined comparing fluorescence signal of samples with known standards.

### Statistics

Group data are presented as mean ± SEM. Statistical comparisons between the two groups were tested using an unpaired t test. Values of p <0.05 were considered statistically significant. All statistical analyses were performed with Prism 8.0.

